# Quartets enable statistically consistent estimation of cell lineage trees under an unbiased error and missingness model

**DOI:** 10.1101/2023.04.04.535437

**Authors:** Yunheng Han, Erin K. Molloy

## Abstract

Cancer progression and treatment can be informed by reconstructing its evolutionary history from tumor cells. However, traditional methods assume the input data are error-free and the output tree is fully resolved. These assumptions are challenged in tumor phylogenetics because single-cell sequencing produces sparse, error-ridden data and because tumors evolve clonally. Here, we find that methods based on quartets (four-leaf, unrooted trees) withstand these barriers. We consider a popular tumor phylogenetics model, in which mutations arise on a (highly unresolved) tree and then (unbiased) errors and missing values are introduced. Quartets are implied by mutations present in two cells and absent from two cells. Our main result is that the most probable quartet identifies the unrooted model tree on four cells. This motivates seeking a tree such that the number of quartets shared between it and the input mutations is maximized. We prove an optimal solution is a consistent estimator of the unrooted cell lineage tree; this guarantee includes the case where the model tree is highly unresolved, with error defined as the number of false negative branches. Lastly, we outline how quartet-based methods might be employed when there are copy number aberrations and other challenges specific to tumor phylogenetics.

## 1 Introduction

Cancer progression and treatment can be informed by reconstructing the evolutionary history of tumor cells (Lim et al. 2020). Although many methods exist to estimate evolutionary trees (called phylogenies) from molecular sequences, traditional approaches assume the input data are error-free and the output tree is fully resolved. These assumptions are challenged in tumor phylogenetics because single-cell sequencing produces sparse, error-ridden data and because tumors evolve clonally so the underlying tree is highly unresolved (Jahn et al. 2016; Schwartz and Schäffer 2017). Here, we demonstrate that quartet-based methods withstand these barriers.

A *quartet* is an unrooted, phylogenetic tree with four leaves. Quartets have long been used as the building blocks for reconstructing the evolutionary history of species (Wilkinson et al. 2005). The reason quartet-based methods have garnered such success in species phylogenetics is their good statistical properties under the Multi-Species Coalescent (`MSC`) model (Pamilo and Nei 1988; Rannala and Yang 2003). An `MSC` model species tree generates gene trees (note that a gene tree reflects the genealogical history of a gene, which is passed down from ancestor to descendant, whereas the species tree governs the pool of potential ancestors). One of the most important results from the last decade of systematics states: for four species, the most probable unrooted gene tree under the `MSC` is topologically equivalent to the unrooted model species tree (Allman et al. 2011). Notably, for trees with more than four leaves, the most probable unrooted gene tree can be topologically discordant with the unrooted model species tree (Degnan 2013). In such situations, the model species tree is said to be in the anomaly zone or the offending gene tree is said to be *anomalous*. It is now widely recognized that anomalous gene trees can challenge traditional species tree estimation methods (Kubatko and Degnan 2007; Roch and Steel 2015).

The statistical theory above has motivated the development of quartet-based methods (e.g., Larget et al. 2010; Mirarab et al. 2014) and is central to their proofs of statistical consistency under the `MSC`. ASTRAL (Mirarab et al. 2014), in particular, has become a gold standard approach to multi-locus species tree estimation. Moreover, new and improved quartet-based methods are continually being developed (Mirarab and Warnow 2015; Zhang et al. 2018; Dibaeinia et al. 2021; Mahbub et al. 2021; Han and Molloy 2023; Zhang and Mirarab 2022). Similar theory and methodology has been given for *triplets*: three-leaf, rooted, phylogenetic trees (Degnan and Rosenberg 2006; Liu et al. 2010; Islam et al. 2020).

Inspired by these efforts, we study the utility of quartets for estimating cell lineage trees under a popular tumor phylogenetics model (Jahn et al. 2016; Ross and Markowetz 2016; Wu 2019; Kizilkale et al. 2022), in which mutations arise on a (highly unresolved) cell lineage tree according to the infinite sites model and then errors and missing values are introduced to the resulting mutation data in an unbiased fashion. The idea is that deviations from a perfect phylogeny can be attributed to sequencing errors, as data produced by single-cell protocols are notoriously error-prone and sparse. Although the infinite sites plus unbiased error and missingness (`IS+UEM`) model generates mutations rather than gene trees, quartets are implied by mutations that are present in two cells and absent from two cells.

Our main result is that there are no anomalous quartets under the `IS+UEM` model; this motivates seeking a cell lineage tree such that the number of quartets shared between it and the input mutations is maximized. We prove an optimal solution to this problem is a consistent estimator of the unrooted cell lineage tree; this guarantee extends to the case of highly unresolved model trees, with error defined as the number of false negative branches. Somewhat surprisingly, our positive results for quartets do not extend to triplets, as there can be anomalous triplets under the `IS+UEM` model under reasonable conditions. The theory presented here generalizes beyond the `IS` model, making it relevant to several recent systematic studies of placental mammals, birds, and bats. Of course, our main focus is tumor phylogenetics, which is additionally challenged by copy number aberrations and doublets. We conclude by outlining how quartet-based methods might be employed given these unique challenges.

## 2. Preliminaries

We now provide some background on phylogenetic trees, models of evolution, and statistical consistency.

### 2.1 Phylogenetic Trees

A *phylogenetic tree* is defined by the triple (*g, X, ϕ*), where *g* is a connected acyclic graph, *X* is a set of labels (often representing species or cells), and ϕ is a bijection between the labels in *X* and leaves (i.e., vertices with degree 1) of *g*. Phylogenetic trees can be either *unrooted* or *rooted*, and we use *u*(*T*) to denote the unrooted version of a rooted tree *T*. Edges in an unrooted tree are undirected, whereas edges in a rooted tree are directed away from the root, a special vertex with in-degree 0 (all other vertices have in-degree 1). Vertices that are neither leaves nor the root are called internal vertices, and edges incident to only internal vertices are called internal edges. An interval vertex with degree greater than 3 (called a *polytomy*) can be introduced to a tree by contracting one of its edges (i.e., deleting the edge and identifying its endpoints). A *refinement* of a polytomy is the opposite of a *contraction*. If there are no polytomies in *T*, we say that *T* is *binary* or *fully resolved* ; otherwise, we say that *T* is *non-binary*. We use the phrase *highly unresolved* to indicate that *T* contains many polytomies and/or that the polytomies in *T* have high degrees.

As previously mentioned, many methods for species tree estimation are based on *quartets*. A quartet is an unrooted, binary tree with four leaves. We denote the three possible quartets on *X* = *{A, B, C, D}* as *q*_1_ = *A, B*|*C, D, q*_2_ = *A, C*|*B, D*, and *q*_3_ = *A, D*|*B, C*. A set of quartets can be created from an unrooted tree *T* by restricting *T* to every possible subset of four leaves (i.e., deleting the other leaves from *T* and then suppressing vertices of degree 2). The resulting set *Q*(*T*) is referred to as the quartet encoding of *T*, and we say that *T* displays quartet *q* if *q ∈ Q*(*T*). Importantly, if *T* contains polytomies, restricting *T* to some subsets of four labels will not produce a quartet. Some selections will produce star trees, which do not provide any topological information. We use *T* |_*S*_ to denote *T* restricted to label set *S* (note that if branch parameters are associated with *T*, they are added together when suppressing vertices of degree 2).

The concepts above for quartets extend to triplets. A *triplet* is a rooted, binary tree with three leaves, and we denote the three possible triplets on *X* = *{A, B, C}* as *t*_*A*_ = *A*|*B, C, t*_*B*_ = *B*|*A, C*, and *t*_*C*_ = *C*|*A, B*. Lastly, a *bipartition* or *split* of label set *X* partitions it into two disjoint subsets. It is easy to see that each edge in an unrooted tree induces a bipartition, and we use *Bip*(*T*) to denote the set of bipartitions induced by all edges in *T*.

### 2.2 Mutations and Models of Evolution

A mutation matrix *M* is an *n × k* matrix, where *n* is the number of rows (representing cells or species) and *k* is the number of columns (representing mutations). Columns are also referred to as characters or site patterns. Our focus here is on 2-state characters, with *M*_*i,j*_ = 0 indicating that mutation *j* is absent from cell *i* and *M*_*i,j*_ = 1 indicating that mutation *j* is present in cell *i*. In tumor phylogenetics, mutations are called in reference to a healthy cell, which is the root of the cell lineage; thus, 0 represents the ancestral state and 1 represents mutant/derived state (note that this interpretation of states 0 and 1 will only be important when looking at triplets and not quartets).

Throughout this paper, we assume the mutation matrix *D* is generated under a *hierarchical model* with two steps (Figure 1).

**Figure 1:**
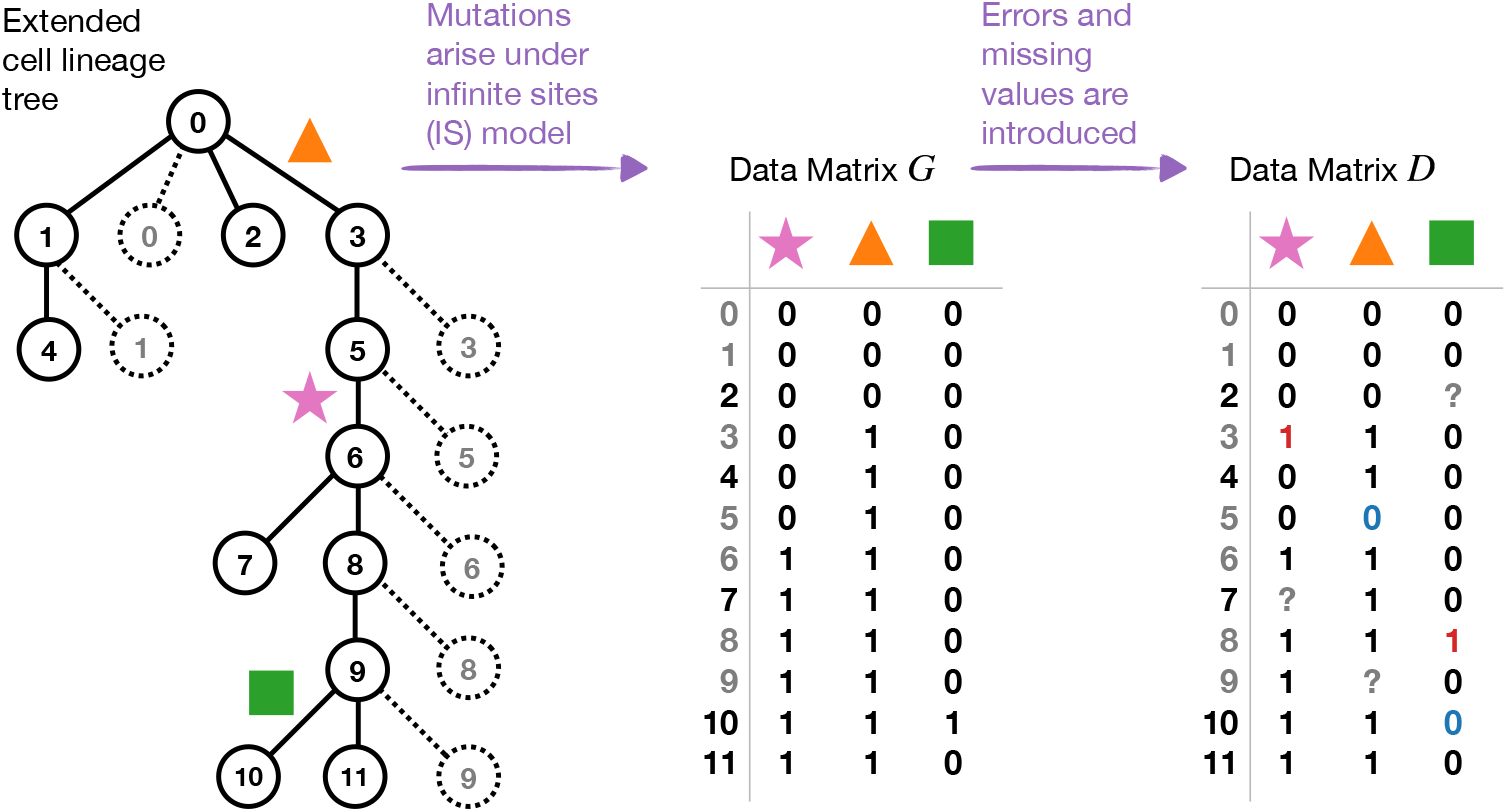
Above we show a model cell lineage tree, where the dashed lines and circles are “fake” edges and vertices, respectively. If we assume a mutation occurs on any non-fake edges with equal probability (as in Kizilkale et al. 2022), then the probability of a mutation on any solid edge will be 1/11. Mutations cannot occur on any of the dashed edges. Data are generated from this model cell lineage tree in two steps. First, mutations arise on the tree under the `IS` model, producing data matrix *G*. Second, false positives (0 flips to 1; shown in red), false negatives (1 flips to 0; shown in blue), and missing values (0/1 flips to ?; shown in grey) are introduced to *G* under the `UEM` model, producing data matrix *D*.

1. A mutation matrix *G* is generated under some model *ℳ*, parameterized by a rooted phylogenetic tree topology *σ* and a set Θ of associated numeric parameters. Importantly, model *ℳ* given (*σ*, Θ) defines a probability distribution on mutation patterns, and we assume mutations in *G* are *independent and identically distributed (i*.*i*.*d*.) according to this model. For simplicity of notation, we typically omit the dependence on the numeric parameters in Θ.

2. Errors and/or missing values are introduced to the ground truth matrix *G* according to the UEM model (described below). This result of this process is the observed matrix *D*.

Hierarchical models of this form, denoted *ℳ*`+UEM`, define a probability distribution on mutation patterns given their parameters. Thus, if we say that *D* is generated under the ℳ+UEM model, then we assume the mutations in *D* are *i*.*i*.*d*. according to this model. We now describe the data generation steps in greater detail for a popular tumor phylogenetics model (Jahn et al. 2016; Ross and Markowetz 2016; Wu 2019; Kizilkale et al. 2022).

#### Step 1: Infinite Sites (IS) model

For tumor phylogenetics, we take *ℳ* to be the infinite sites (`IS`) model, so the mutation matrix *G* is generated under the `IS` model given a rooted cell lineage tree *σ* and a set Θ of edge probabilities that sum to 1. Specifically, every edge *e* in *σ* is associated with a numeric value *p*(*e*) *∈* Θ, indicating the probability that a mutation occurs on *e*. When a mutation occurs on *e*, all cells on a directed path from *e* to any of the leaves of *σ* are set to state 1; all other cells are set to state 0. Thus, a mutation corresponds to the bipartition induced by the branch on which it occurred. Edges on which mutations cannot occur are contracted so that the probability of a mutation on any edge in *σ* is strictly greater than zero. An exception occurs because the model cell lineage tree must be “extended” so that each internal vertex is also represented as a “fake” leaf. Mutations are not allowed on “fake” edges, which are incident to “fake” leaves. This extension process is necessary because single-cell sequencing can produce data for cells that are ancestral to other cells in the same data set.

#### Step 2: Unbiased Error and Missingness (UEM) model

If mutation matrix *G* is generated under the IS model given (*σ*, Θ), then reconstructing *σ* is trivial. However, for tumor phylogenetics, false positives and false negatives are introduced to *G*, producing the observed matrix *D*. This is done according to

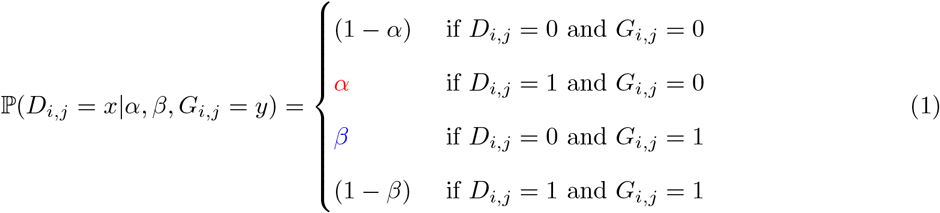

where 0 ≤ *α* < 1 and 0 ≤ *β* < 1 are the probability of false positives and false negatives, respectively. Simultaneously, missing values are introduced to *G* with probability 0 ≤ *γ* < 1; this can be incorporated into the model by multiplying each of the cases in Equation 1 by (1 –*γ*).

Our goal is to estimate cell lineage trees under the `IS+UEM` model. An important property for phylogeny estimation methods is whether they are statistically consistent under the model of interest.

##### Definition 1.

*[Statistical Consistency; see Section 1.1 of Warnow 2017] Let 𝒜 be some model that generates mutations, and let D be a mutation matrix, with n rows (cells or species) and k columns (mutations), generated under 𝒜 given rooted tree σ and numerical parameters* Θ. *We say that an estimation method is statistically consistent under 𝒜 if for any ϵ >* 0, *there exists a constant K >* 0 *such that when D contains at least K mutations, the method given D returns (the unrooted version of) σ with probability at least* 1 – *ϵ. Alternatively, we might say that the error in the tree estimated from D is zero with probability at least* 1 – *ϵ*.

The idea is that as the number *k* of mutations goes towards infinity, the error in the estimated tree is zero with high probability. Tree error is typically defined as the number of *false negative branches* (i.e., branches in *σ* that are missing from the estimated tree) plus the number of *false positive branches* (i.e., branches in the estimated tree that are missing from *σ*).

## 3 No anomalous quartets under an unbiased error and missingness model

To begin, we assume that the rooted cell lineage tree *σ* has four leaves; therefore, it must have one of five tree shapes shown in Figure 2. Two of them display a star when unrooted, and the other three correspond to a quartet when unrooted.

**Figure 2:**
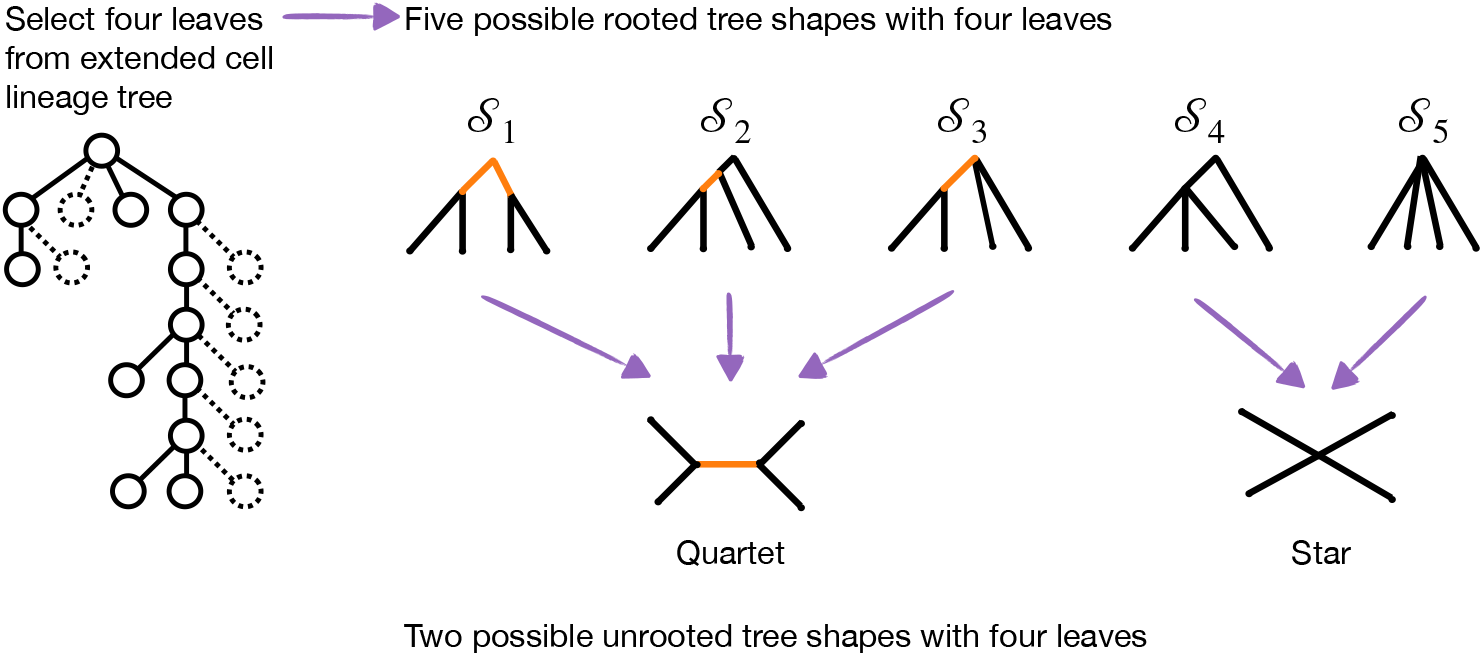
There are five possible tree shapes with four leaves. Three of these tree shapes (*S*_1_, *S*_2_, and *S*_3_) have a non-trivial unrooted topology (quartet). The other two tree shapes (*S*_4_ and *S*_5_) have a trivial unrooted topology (star). Looking at Figure 1, tree shapes *S*_1_–*S*_5_ are observed by sampling cells *X*_1_ = {1, 4, 10, 11*}, X*_2_ = {1, 7, 10, 11}, *X*_3_ = {1, 2, 10, 11}, *X*_4_ = {1, 9, 10, 11}, and *X*_5_ = {0, 1, 2, 3}, respectively.

If mutations are generated *i*.*i*.*d*. under some model *A* given *σ*, there are 16 possible patterns on four cells, denoted *{A, B, C, D}*. A quartet is implied by two cells being in state 1 and two cells being in state 0. Therefore, two patterns (*ABCD* = 0011 and 1100) support quartet *q*_1_ = *A, B*|*C, D*, two patterns (0101 and 1010) support quartet *q*_2_ = *A, C*|*B, D*, two patterns (0110 and 1001) support quartet *q*_3_ = *A, D*|*B, C*, and the other 10 patterns do not provide topological information. Henceforth, we denote the probability of quartets under model *A* given *σ* as 𝕡_*𝒜*_(*q*_1_|*σ*) = 𝕡_*𝒜*_(1100|*σ*) + 𝕡_*𝒜*_(0011|*σ*), 𝕡_*𝒜*_(*q*_2_|*σ*) = 𝕡_*𝒜*_(1010|*σ*) + 𝕡_*𝒜*_(0101|*σ*), and 𝕡_*𝒜*_(*q*_3_|*σ*) = 𝕡_*𝒜*_(1001|*σ*) + 𝕡_*𝒜*_(0110|*σ*). Now we consider quartet-informative patterns generated from a model tree with more than four leaves.

### Definition 2

(No anomalous quartets). *We say that there are no anomalous quartets under model 𝒜 given rooted tree σ if the following inequalities hold for every subset S of four species in σ. Let q*_1_, *q*_2_, *q*_3_ *denote the three quartets on S, and let i index {*1, 2, 3}.

1. *If u*(*σ*)|_*S*_ = *q*_*i*_, 𝕡 _*𝒜*_(*q*_*i*_|*σ*) *>* 𝕡 _*𝒜*_ (*q*_*j*_|*σ*) *for all j ∈ {*1, 2, 3*} such that i* ≠ *j*.
2. *If u*(*σ*)|_*S*_ *is a star*, P_*𝒜*_(*q*_1_|*σ*) = P_*𝒜*_ (*q*_2_|*σ*) = P_*𝒜*_(*q*_3_|*σ*).

This brings us to the main result of this section.

### Theorem 1.

*There are no anomalous quartets under the IS+UEM model, assuming α* + *β ≠* 1.

The statement above directly follows from Lemma 1 and Corollary 1.

### Lemma 1.

*There are no anomalous quartets under the* `IS` *model. Moreover, all quartet-informative patterns have zero probability except for one or both of the patterns corresponding to u*(*σ*) *when u*(*σ*) *is not a star*.

If *σ* has more than four leaves, we can restrict *σ* to any subset of four leaves and get a valid sub-model (i.e., a sub-model for which the probability of the mutation patterns on four cells is the same as under the larger model tree). The sub-model is formed by deleting the other leaves and adding branch parameters together when suppressing vertices of degree 2. The mutation pattern probabilities for the four cells under this sub-model will be the same as the larger tree because addition represents an *or* condition (i.e., a mutation occurring on this branch *or* on that branch will produce the same pattern when looking at only a subset of cells). Thus, it suffices to verify that there are no anomalous quartets for *σ* with four leaves. This can be done by considering a mutation occurring on each of the internal branches of all possible rooted tree shapes with four leaves (Figure 2) and comparing the resulting pattern to the unrooted tree shape; see Supplementary Materials for details.

The following two lemmas will also be useful later.

### Lemma 2.

*Let* 0 ≤ *α* < 1 *and* 0 ≤ *β* < 1. *Then*,

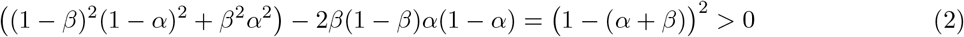

*for α* + *β ≠* 1. *If α* + *β* = 1, *the inequality in Equation 2 becomes an equality*.

The statement above follows from expanding the polynomials; see the Supplementary Materials for details.

### Lemma 3.

*If there are no anomalous quartets under model M, then there are no anomalous quartets under the ℳ+* `UE` *model, assuming α* + *β ≠* 1.

*Proof*. Taking any subset of four leaves, there are 16 possible mutation patterns that may occur under model *M*. These are the two invariant patterns (0000 and 1111), the eight variant but quartet-uninformative patterns (1000, 0100, 0010, 0001, 0111, 1011, 1101, 1110), and the six quartet-informative patterns (1100, 0011, 0101, 1010, 0110, 1001). For each pattern *g* listed above, we enumerate all possible ways of introducing errors (false positives and false negatives); this gives us the probability of each of the 16 mutation patterns under the UE model given (*α, β*). Now we need to put this information together to get the probability of quartets under the *ℳ* `+UE` model. First, we compute the probability of observing any quartet *q* from errors (false positives and false negatives) being introduced to the invariant and variant but quartet-uninformative characters

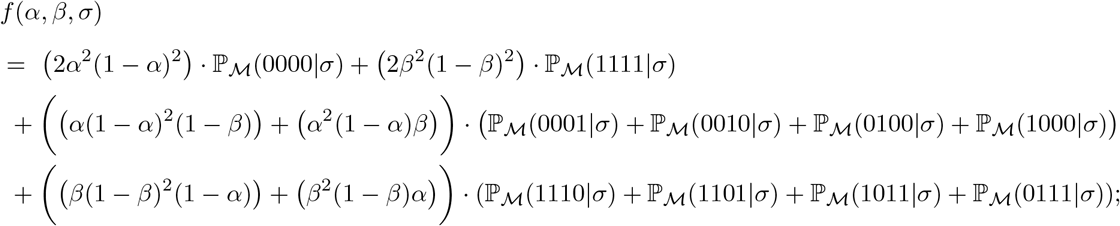

see Supplementary Tables S1–S4 for details. Second, we repeat this calculation for the quartet-informative patterns; see Table 1 and Supplementary Tables S5–S6 for details. Putting it all together gives us the probability of each quartet under the ℳ `+UE` model

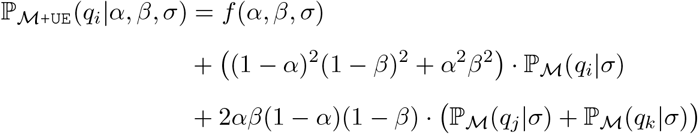

for *i, j, k* ∈ {1, 2, 3} such that *i* ≠ *j* ≠ *k*. Now we can compute the difference in probabilities between quartets *q*_*i*_ and *q*_*j*_ under the *ℳ* `+UE` model for any *i, j* ∈ {1, 2, 3} such that *i* ≠ *j*. By Lemma 1, we have

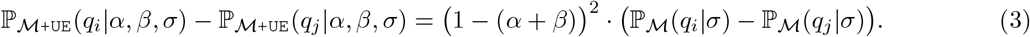

**Table 1:**
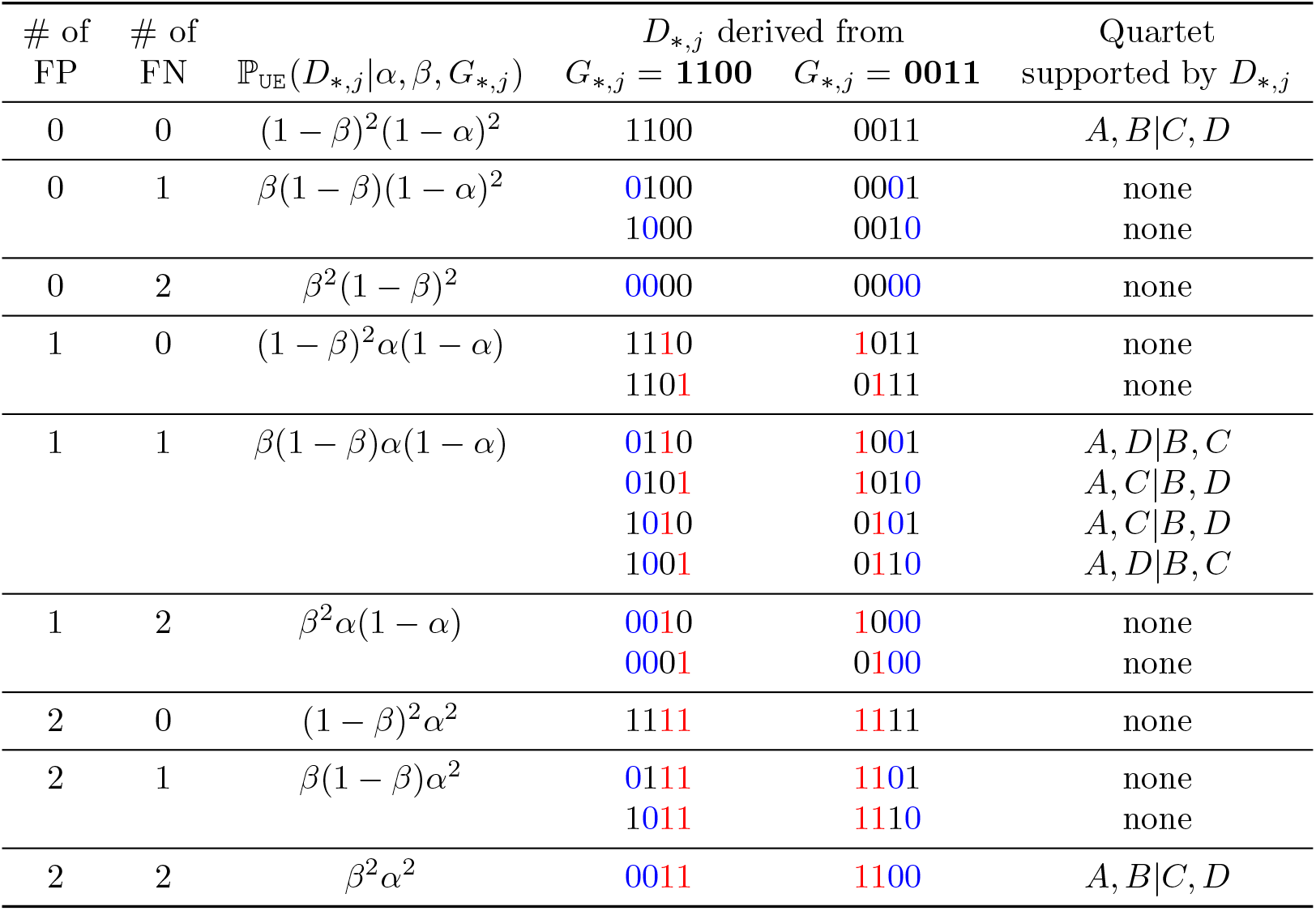
Above we show the mutation patterns with four cells *{A, B, C, D}* that can be generated under the `UE` model given **pattern #12** (*G*_*\*,j*_ = 1100) and **pattern #3** (*G*_*\*,j*_ = 0011). The red values indicate a false positive introduced to *G*_*\*,j*_ by flipping 0 to 1. The blue values indicate a false negative introduced to *G*_*\*,j*_ by flipping 1 to 0. Similar tables for the other 14 patterns are provided in the Supplementary Materials.

Assuming *α* + *β* ≠ 1, this quantity is zero if 𝕡_*ℳ*_(*q*_*i*_|*σ*) = 𝕡_*ℳ*_ (*q*_*j*_|*σ*) and greater than zero if 𝕡_*ℳ*_ (*q*_*i*_|*σ*) *>* 𝕡_*ℳ*_ (*q*_*j*_|*σ*). Because there are no anomalous quartets under model *M*, the former will be the case if *𝒰* (*σ*) is a star; the latter will be the case if *u*(*σ*) = *q*_*i*_. It follows there are no anomalous quartets under the *M*`+UE` model.

Note that the quantity *α* + *β* is unlikely to equal 1 in practice, as both probabilities should be less than 0.5. We now extend the result above to address unbiased missing values.

### Corollary 1.

*If there are no anomalous quartets under model ℳ, then there are no anomalous quartets under the ℳ+UEM model, assuming that α* + *β ≠* 1.

*Proof*. If one or more of the values in a mutation pattern is missing, then no quartet is displayed. Thus, unbiased missingness can be accounted for in the proof of Lemma 3 simply by updating 𝕡_*ℳ*_ (*x*|*σ*) to (1 *− γ*)^4^ · 𝕡_*ℳ*_ (*x*|*σ*). In this case, Equation 3 becomes

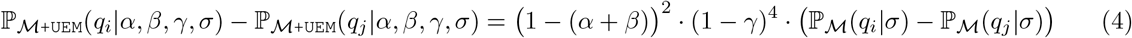

which does not change our argument.

This concludes the derivation of the main result of this section (Theorem 1). Before moving on to triplets, we note that the difference in quartet probabilities (Equation 4) depends on (1) the probability of false positives and negatives (specifically how close their sum is to one), (2) the probability of missing values, and (3) the probability of observing quartet-informative patterns under model *ℳ* given *σ*. In the simulations performed by Kizilkale et al. (2022), the largest values of *α, β*, and *γ* were 0.001, 0.2, and 0.05, respectively. In this scenario with *u*(*σ*) = *q*_*i*_, we have 𝕡_IS+UEM_(*q*_*i*_|*α, β, γ, σ*) *−* 𝕡_IS+UEM_(*q*_*j*_|*α, β, γ, σ*) = 0.52 · 𝕡_IS_(*q*_*i*_|*σ*) for any *i, j ∈ {*1, 2, 3} such that *i* ≠ *j*. The magnitude of P_IS_(*q*_*i*_|*σ*) depends on the lineage tree and the four cells sampled from it. We discuss sampling further in the context of triplets; for now, we note that in practical settings, 𝕡_IS_(*q*_*i*_|*σ*) may have a greater impact on the difference in quartet probabilities under the IS+UEM model than *α, β*, or *γ*.

## 4 Anomalous triplets under an unbiased error and missingness model

We now derive related results for triplets. To begin, we assume the rooted cell lineage tree *σ* has three leaves; therefore, it must have one of two topologies: binary or non-binary (Supplementary Figure S1). If mutations are generated *i*.*i*.*d*. under some model *A* given *σ*, there are 8 possible patterns on three cells, denoted *{A, B, C}*. A triplet is implied by two cells being in state 1 (i.e., the mutant/derived state) and one cell being in state 0 (i.e., the ancestral state). One pattern (*ABC* = 110) supports triplet *t*_*C*_ = *C*|*A, B*, one pattern (101) supports triplet *t*_*B*_ = *B*|*A, C*, one pattern (011) supports triplet *t*_*A*_ = *A*|*B, C*, and the other five patterns do not provide topological information. Henceforth, we denote the probability of triplets under model *A* given *σ* as 𝕡_*ℳ*_ (*t*_*A*_|*σ*) = 𝕡_*ℳ*_ (011|*σ*), 𝕡_*ℳ*_ (*t*_*B*_|*σ*) = 𝕡_*ℳ*_ (101|*σ*), and 𝕡_*ℳ*_ (*t*_*C*_|*σ*) = 𝕡_*ℳ*_ (110|*σ*). Now we consider triplet-informative patterns generated from a model tree with more than three leaves.

### Definition 3

(No anomalous triplets). *We say that there are no anomalous triplets under model A if the following inequalities hold for every subset S of three species in σ. Let t*_1_, *t*_2_, *t*_3_ *denote the three triplets on S, and let i index {1, 2, 3}*.

*1. If σ*|_*S*_ = *t*_*i*_, 𝕡_*ℳ*_ (*t*_*i*_|*σ*) *>* 𝕡_*ℳ*_ (*t*_*j*_|*σ*) *for all j ∈ {*1, 2, 3*} such that i* ≠ *j*.

*2. If σ*|_*S*_ *is non-binary*, P_*A*_(*t*_1_|*σ*) = P_*A*_(*t*_2_|*σ*) = P_*A*_(*t*_3_|*σ*).

This brings us to the main result of this section.

### Theorem 2.

*There are no anomalous triplets under the IS+UEM model, assuming one of two conditions:*

*(1) α* = 0 *or (2) α* + *β* ≠ 1 *and* 𝕡_*IS*_(100|*σ*) = P_*IS*_(010|*σ*) = 𝕡_*IS*_(001|*σ*). *Otherwise, there can be anomalous triplets under the IS+UEM model*.

The statement above directly follows from Lemma 4 and Corollary 2.

### Lemma 4.

*There are no anomalous triplets under the* `IS` *model. Moreover, all triplet-informative patterns have zero probability except for the pattern corresponding to σ when σ is not non-binary*.

If *σ* has more than three leaves, we can restrict *σ* to any subset of three leaves and get a valid sub-model (i.e., a sub-model for which the probability of the mutation patterns on three cells is the same as under the larger model tree, as discussed for quartets). Thus, it suffices to verify that there are no anomalous triplets for *σ* with three leaves. This can be done by considering a mutation occurring on each of the internal branches of all possible rooted tree shapes with three leaves (Supplementary Figure S1) and comparing the resulting pattern to the tree shape; see Supplementary Materials for details.

### Lemma 5.

*If there are no anomalous triplets under model ℳ, then there are no anomalous triplets under the ℳ*`+UE` *model, assuming one of two conditions: (1) α* = 0 *or (2) α* + *β* ≠ 1 *and* 𝕡_*ℳ*_(100|*σ*) = 𝕡_*ℳ*_ (010|*σ*) =𝕡_*ℳ*_ (001|*σ*). *Otherwise, there can be anomalous triplets under the ℳ* `+UE` *model*.

*Proof*. Taking any subset of three leaves, there are 8 possible mutation patterns that may occur under model ℳ. These are the two invariant patterns (*ABC* = 000 and 111), the three variant but triplet-uninformative patterns (100, 010, 001), and the three triplet-informative patterns (110, 101, 011). For each pattern *g* listed above, we enumerate all possible ways of introducing errors (false positives and false negatives); this gives us the probability of mutation patterns under the `UE` model given (*α, β, g*); see Supplementary Tables S7–S14. Putting everything together, we find that the probability of triplet *t*_*C*_ under the *ℳ*+UE model is

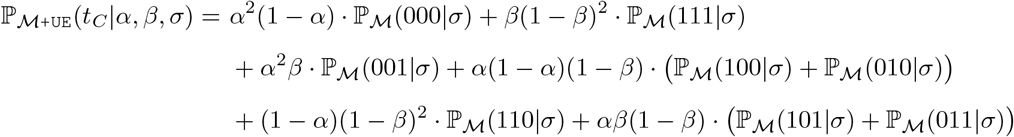

Similar probabilities can be computed for *t*_*B*_ and *t*_*A*_. To provide a general formula, we define

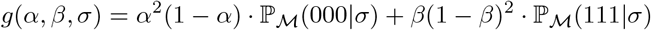

and set *x*_*A*_ = 100, *x*_*B*_ = 010, and *x*_*C*_ = 001. This allows us to write the probability of any triplet as

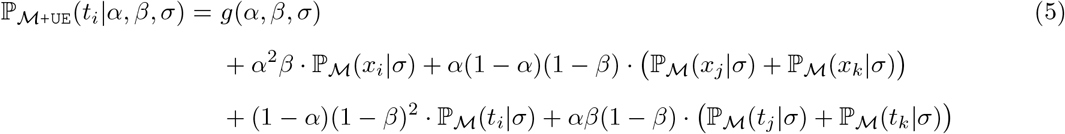

where *i, j, k ∈ {A, B, C}* with *i ≠ j ≠ k*. Now we can compute the differences in probabilities between *t*_*i*_and *t*_*j*_ under the *ℳ* `+UE` for any *i, j ∈ {A, B, C}* such that *i ≠ j*. We find

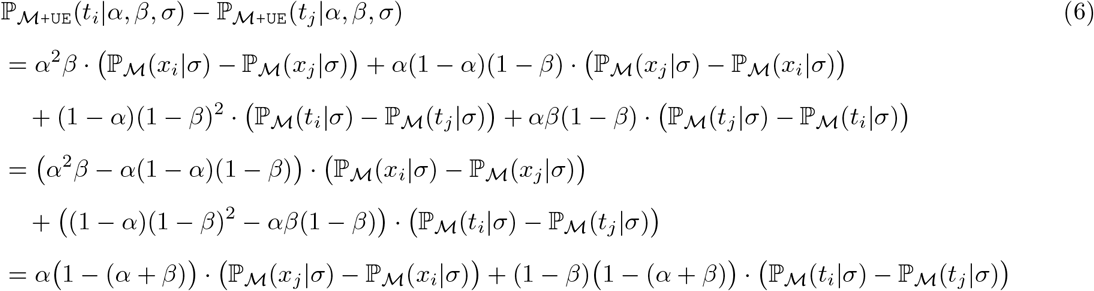

Assuming that either condition (1) *α* = 0 or condition (2) *α* + *β* ≠ 0 and 𝕡_*ℳ*_ (*x*_*i*_|*σ*) = 𝕡_*ℳ*_ (*x*_*j*_|*σ*), this quantity is zero if 𝕡_*ℳ*_ (*t*_*i*_|*σ*) = 𝕡_*ℳ*_ (*t*_*j*_|*σ*) and is greater than zero if 𝕡_*ℳ*_ (*t*_*i*_|*σ*) *>* 𝕡_*ℳ*_ (*t*_*j*_|*σ*). Because there are no anomalous triplets under *M*, the former will be the case if *σ* is non-binary; the latter will be the case if *σ* = *t*_*i*_. It follows there are no anomalous triplets under the *M*+UE model, provided one of the two conditions hold. If these conditions do not hold and *σ* ≠ *t*_*j*_, triplet *t*_*j*_ is anomalous when

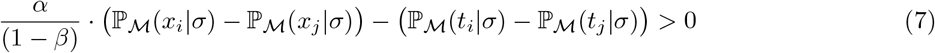

for *i, j ∈ {A, B, C}* such that *i* ≠*j*.

The result above extends easily to the case with unbiased missing values (see Corollary 1).

### Corollary 2.

*If there are no anomalous triplets under model M, then there are no anomalous triplets under the M+UEM model, assuming that one of the two conditions: (1) α* = 0 *or (2) α* + *β ≠* 1 *and* 𝕡_*ℳ*_ (100|*σ*) = 𝕡_*ℳ*_ (010|*σ*) = 𝕡_*ℳ*_ (001|*σ*). *Otherwise, there can be anomalous triplets under the M+UEM model*.

This concludes the derivation of the main result of this section (Theorem 2). Before moving onto phylogeny estimation, we consider whether anomalous triplets should be expected in the context of tumor phylogenetics, so setting *M* to be the `IS` model. As previously mentioned, in the simulations performed by Kizilkale et al. (2022), the largest values of *α* and *β* were 0.001 and 0.2, respectively. This would put *α/*(1 *− β*) = 0.00125 in Equation 7, so 𝕡_*ℳ*_ (*x*_*i*_|*σ*) *−* 𝕡_*ℳ*_ (*x*_*j*_|*σ*) would need to be 800 times greater than 𝕡_*ℳ*_ (*t*_*i*_|*σ*) *−* 𝕡_*ℳ*_ (*t*_*j*_|*σ*) for *t*_*j*_ to be anomalous under the *M*+UEM model (note the terms for missing values will cancel out in Equation 7). Although this seems drastic, it could occur when *σ* is created by restricting a larger model tree to a subset three leaves. Consider the cell lineage tree in Figure 1 but with 1000 additional cells added between cell 9 and cell 10 (recall mutations occur on all non-fake edges with equal probability in this example). If we sample cells {1, 4, 10}, the resulting sub-model *σ* has rooted topology *t*_10_ = 10|1, 4 and defines the following probability distribution on mutations. For the variant but triplet-uninformative patterns, we have 𝕡_IS_(*x*_1_|*σ*) = 0, 𝕡_IS_(*x*_4_|*σ*) = 1*/*1012, and 𝕡_IS_(*x*_10_|*σ*) = 1009*/*1012, and for triplet-informative patterns, we have P_IS_(*t*_1_|*σ*) = 0, 𝕡_IS_(*t*_4_|*σ*) = 0, and 𝕡_IS_(*t*_10_|*σ*) = 1*/*1012. Now looking at Equation 7, we find that triplets *t*_1_ and *t*_4_ are anomalous under the `IS+UEM` model. Based on this analysis, we conjecture that triplets are less robust to error than quartets when sampling cells from a larger cell lineage tree.

## 5. Phylogeny Estimation from Quartets

Because there are no anomalous quartets under the `IS+UEM` model under reasonable assumptions (i.e., *α* + *β* ≠ 1), we now consider the utility of quartet-based methods for estimating cell lineage trees from mutation data. By quartet-based methods, we mean heuristics for the Maximum Quartet Support Supertree (MQSS) problem (Wilkinson et al. 2005; also see Section 7.7 in Warnow 2017).

### Definition 4

(Maximum Quartet Support Supertrees). *The MQSS problem is defined by*

***Input:*** *A set of unrooted trees P* = *{T*_1_, *T*_2_, …, *T*_*k*_*}, with tree T*_*i*_ *on leaf label set X*_*i*_

***Output:*** *A unrooted, binary tree B on leaf label set* 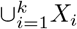 *that maximizes QS*_*D*_(*B*) =∑_*q∈Q*(*B*)_ *w*_*P*_ (*q*), *where w*_*P*_ (*q*) *is the number of input trees in 𝒫 that display q*

When all input trees are on the same leaf label set, MQSS becomes weighted quartet consensus (WQC). An optimal solution to WQC is a consistent estimator of the unrooted species tree topology under the MSC model (Theorem 2 in Mirarab et al. 2014). Although MQSS and WQC are NP-hard (Jiang et al. 2001; Lafond and Scornavacca 2019), fast and accurate heuristics have been developed, with the most well-known being ASTRAL (Mirarab et al. 2014). Since version 2 (Mirarab and Warnow 2015; Zhang et al. 2018), ASTRAL allows the input trees to be incomplete; it is statistically consistent under the MSC model, provided some assumptions on missing data (see Nute et al. 2018 for details).

There are two notable differences when using quartet-based methods to reconstruct cell lineage trees, rather than species trees. The first difference is that input are mutations rather than unrooted (gene) trees. This issue was addressed by Springer et al. (2019), who treat mutations as unrooted trees with at most one internal branch (Figure 3). Given this transformation of the input, it is possible to run ASTRAL and other quartet-based methods on mutation data. The second difference is that ASTRAL outputs a binary tree; however, the model cell lineage tree is unlikely to be binary given the clonal model of tumor evolution (see review by Schwartz and Schäffer 2017). Indeed, methods for tumor phylogenetics are often evaluated on data simulated from highly unresolved trees (Kizilkale et al. 2022). Our next result suggests that MQSS/WQC are useful problem formulations, even when data are generated from non-binary trees.

**Figure 3:**
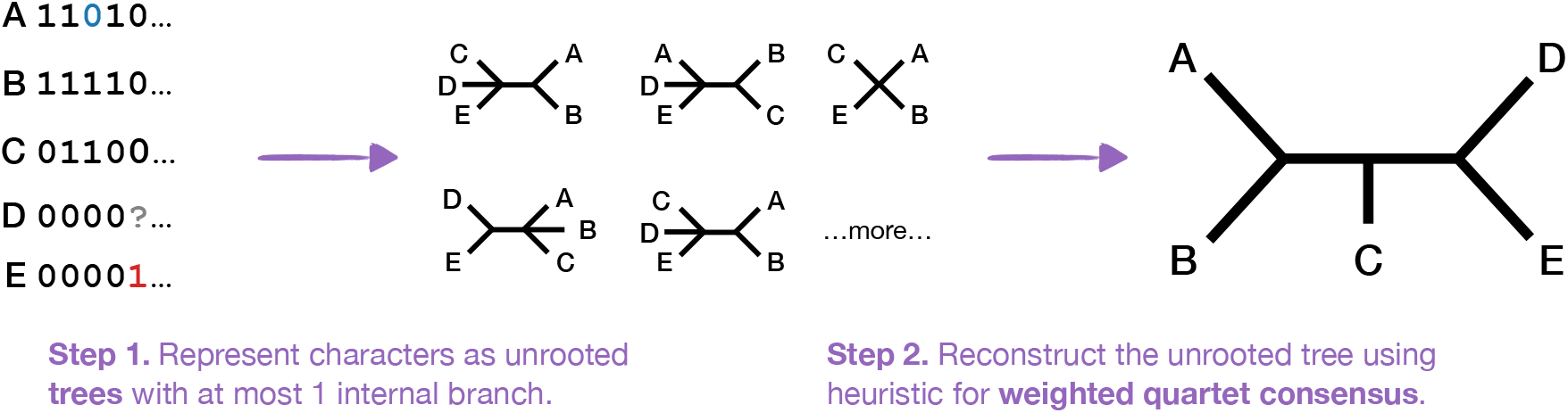
This figure shows a mutation matrix with one false negative (in blue), one false positive (in red), and one missing entry. The first step indicates how each mutation (column in the matrix) corresponds to an unrooted tree with at most one internal branch. The second step is to estimate the phylogeny by applying a quartet-based method. This approach is motivated by there being no anomalous quartets under the `IS+UEM` model, assuming *α* + *β* ≠ 1 (Theorem 1).

### Theorem 3.

*Let σ be a rooted tree with at least one internal branch when unrooted (so it can be non-binary), and let D be an n × k mutation matrix generated under the* `IS+UEM` *model given* (*α, β, γ, σ*). *Suppose that α* + *β ≠* 1. *Then, an optimal solution to MQSS given D is a consistent estimator of u*(*σ*) *under the IS+UEM model, with tree error defined as the number of false negative branches. If in addition* 0 *< α, β, <* 1, *ASTRAL given D is statistically consistent under the* `IS+UEM` *model, with tree error defined as the number of false negative branches*.

The first statement above follows from Theorem 1 and Lemma 6. The second statement follows from Theorem 1, Corollary 3, and the observation that every complete mutation pattern is possible under the `IS+UEM` model when 0 *< α, β <* 1.

### Lemma 6.

*Let σ be a rooted tree with at least one internal branch when unrooted (so it can be non-binary), and let D be an n × k mutation matrix generated under model A given σ. If there are no anomalous quartets under A, an optimal solution to MQSS given D is a consistent estimator of u*(*σ*) *under model A, with tree error defined as the number of false negative branches*.

*Proof*. Let *B* be an unrooted, binary tree on the same label set as *𝒰* (*σ*). The number of false negative branches between *B* and *𝒰* (*σ*) is zero if *B* is a refinement of *u*(*σ*), meaning that *B* can be obtained from *𝒰* (*σ*) in a sequence of refinement operations (this sequence has length zero if *σ* is binary). Thus, to prove consistency with tree error defined as the number of false negative branches, we revise Definition 1 to say that for any *ϵ>* 0, there exists a constant *K >* 0 such that when *D* contains at least *K* mutations, an optimal solution to MQSS given *D* is a refinement of *𝒰* (*σ*) with probability at least 1 *− E*. The remainder of the proof follows from Lemma 7.

### Lemma 7.

*Suppose the conditions of Lemma 6 hold. Let L*(*σ*) *be the label set of σ, and let B and T be unrooted, binary trees on L*(*σ*). *Suppose that B is a refinement of 𝒰* (*σ*) *and that T is NOT. Then, for any pair B and T and for any E >* 0, *there exists a constant K >* 0 *such that when D contains at least K mutations, QS*_*D*_(*B*) *> QS*_*D*_(*T*) *with probability at least* 1 *− ϵ*.

*Proof*. To begin, we restate the inequality as

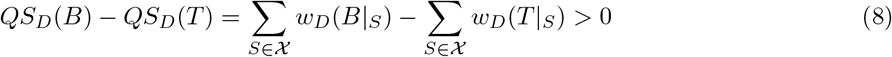

where *𝒳* is the set of all possible subsets of four elements from *L*(*σ*) and *w*_*D*_(*q*) is the number of mutations in *D* that imply quartet *q*.

### Claim 1

First, we claim that as *k → ∞, w*_*D*_*/k* converges to its expectation *F* ^*\**^ under model *A* given *σ* with probability 1. Claim 1 holds by the strong law of large numbers, as noted in the proofs of consistency for quartet and triplet-based methods under the MSC model (see Liu et al. 2010 for an example). Let 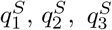 denote the three possible quartets on *S*. Then, we can re-state Claim 1 as follows. As 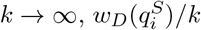 converges to 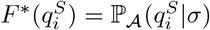 with probability 1 for all *i ∈ {*1, 2, 3} and for all *S ∈ 𝒳*.

### Claim 2

Second, we claim there exists a *δ* such that whenever ∥*w*_*D*_*/k − F*_*\**_∥_*∞*_ *< δ*, Equation 8 holds.

We show Claim 2 is true for *δ* = *π/*2| *𝒳* |, where

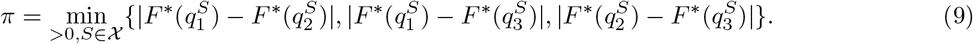

There are three cases to consider.

*Case 1:* Let *𝒲 ⊂ 𝒳* include all *S ∈ 𝒳* such that *B*|_*S*_ and *T* |_*S*_ are the same quartet. Then, *w*_*D*_(*B*|_*S*_) = *w*_*D*_(*T* |_*S*_) for all *S ∈ 𝒲*, giving us

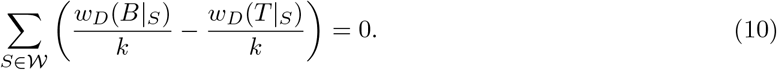

Now suppose that *B* and *T* restricted to *S* display different quartets, denoted 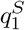 and 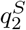, respectively. If 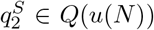, it must also be in *Q*(*B*) as *Q*(*u*(*N*)) *⊂ Q*(*B*). Therefore, *q*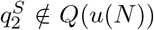. This gives us two additional cases to consider.

*Case 2:* Let *Y ⊂ X* include all *S ∈ 𝒳* such that *B*|_*S*_ and *T* |_*S*_ are different quartets, denoted 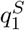 and 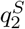, respectively, with 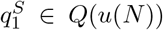 and 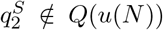. Then, for all 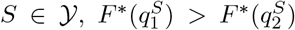 and 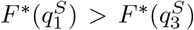 because there are no anomalous quartets under *A* (Definition 2). Therefore, whenever ∥*w*_*D*_*/k − F*_*\**_ ∥ _*∞*_ *< π/*2, *w*_*D*_(*B*|_*S*_) *> w*_*D*_(*T* |_*S*_) for all *S ∈ 𝒴*, giving us

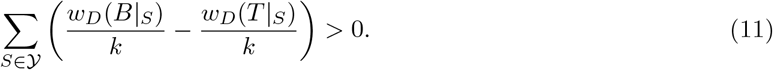

*Case 3 (only needed for non-binary σ):* Let *Z ⊂ 𝒳* include all *S ∈ 𝒳* such that *B*|_*S*_ and *T* |_*S*_ are different quartets, denoted 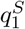 and 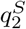, respectively, with 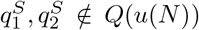. By our assumptions on *B, N* |_*S*_ is a star. Then, because there are no anomalous quartets under *A* (Definition 2), 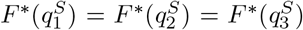. However, even as *k → ∞*, we are not guaranteed to get an exact equality 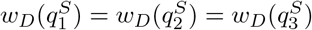. Thus, we need to put an upper bound *δ* on *∥ w*_*D*_*/k −F*_*\**_ *∥* _*∞*_ so that Equation 8 holds even when *w*_*D*_(*B*|_*S*_) *< w*_*D*_(*T* |_*S*_) for all *S ∈ Z*. This happens for *δ* = *π/*2|*X* |. When *∥ w*_*D*_*/k − F*_*\**_ ∥ _*∞*_ *< π/*2|*X* |, we have

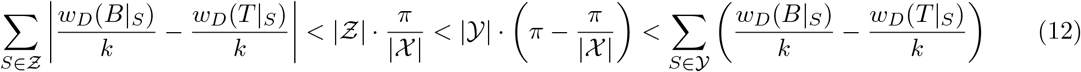

because (𝒴 + *𝒵*)*/ 𝒳 ≤* 1 and 1 *≤* | *𝒴* |. If not, either *T* is a refinement of *𝒰* (*σ*) or *𝒰* (*σ*) has no internal branches, contradicting our assumptions.

*Putting the cases together:* If *𝒰* (*σ*) is binary, then *𝒲* | *𝒴* is a partition *𝒳*, so we can combine equations 10 and 11 to get Equation 8. If *𝒰* (*σ*) is non-binary, then *𝒲* | *𝒴* |*Z* is a partition of *𝒳*, so we can combine Equations 10 and 12 to get Equation 8.

**Wrap up**. By Claim 1, for any *ϵ >* 0, there exists a constant *K >* 0 such that when *D* contains at least *K* mutations, ‖ *w*_*D*_*/k − F* ^*\**^ ‖ _*∞*_ *< π/*2|*X* | with probability at least 1 *− E*. Then, by Claim 2, for any *E >* 0, when *D* contains at least *K* mutations, *QS*_*D*_(*B*) *> QS*_*D*_(*T*) with probability at least 1 *− ϵ*.

### Corollary 3

*Suppose the conditions of Lemma 6 hold. If there are no anomalous quartets under model 𝒜 and every complete mutation pattern occurs with non-zero probability under 𝒜, then ASTRAL given 𝒟 is statistically consistent under 𝒜, with tree error defined as the number of false negative branches*.

*Proof*. ASTRAL solves MQSS exactly within a constrained version of the solution space, denoted σ. Its algorithm has two main steps: first, set σ so that it contains all bipartitions induced by the input “trees” (i.e., mutations in *D*), and second find a solution *B* to MQSS such that *Bip*(*B*) *⊆* σ. Because all complete mutation patterns have non-zero probability under the model *A*, for any *ϵ* _1_ *>* 0, there exists a constant *K*_1_ *>* 0 such that when *k ≥ K*_1_, σ will contain all bipartitions induced by at least one refinement of *𝓊* (*σ*) with probability 1 *− E*_1_. By Lemma 7, for any *ϵ* _2_ *>* 0, there exists a constant *ϵ* _2_ *>* 0 such that when *k ≥ K*_2_, *QS*_*D*_(*B*) *> QS*_*D*_(*T*) for any pair *B* and *T* of unrooted binary trees on *L*(*σ*) such that *B* is a refinement of 𝓊 (*σ*) and *T* is NOT. Now let *E >* 0 and select *ϵ* _1_, *ϵ* _2_ *>* 0 such that *ϵ* _1_ + *ϵ* _2_ *< E*. Then, when *D* contains at least max{ *ϵ* _1_, *ϵ* _2_} mutations, ASTRAL given *D* returns a refinement of *𝓊* (*σ*) with probability at least (1 *− ϵ* _1_)(1 *− ϵ* _2_) *>* (1 *− ϵ*). It follows the number of false negative branches is zero with probability at least 1 *− ϵ*.

Lastly, we note that related results can be derived for triplets by viewing mutations as a rooted trees with at most one internal branch (Supplementary Figure S2).

### Theorem 4

*Suppose that the conditions of Theorem 3 hold but that α* = 0 *(instead of α* + *β* ≠1*). Then, an optimal solution to Maximum Triplet Support Supertree (MTSS) problem given D is a consistent estimator of σ under the* `IS+UEM` *model, with tree error defined as the number of false negative branches*.

This result follows from Theorem 2 and Lemma 6 but replacing “quartet” with “triplet” and “rooted” with “unrooted”.

## 6. Discussion

Quartet-based approaches have garnered much success for estimating species phylogenies under the Multi-Species Coalescent (Mirarab et al. 2014; Mirarab and Warnow 2015; Zhang et al. 2018). Here, we considered their application for estimating cell lineage trees, focusing on two important differences between estimating cell lineage trees compared to species trees. First, errors and missing values can arise from single-cell sequencing and thus are typically modeled. Second, the model cell lineage tree may be highly unresolved because tumors evolve clonally. To address these issues, we first show that there are no anomalous quartets under the infinite sites (`IS`) plus unbiased error and missingness (`UEM`) model, which is widely used in tumor phylogenetics (this is an *identifiability result*). We then show that under the `IS+UEM` model, an optimal solution to the Maximum Quartet Support Supertree (MQSS) problem is a refinement of the model cell lineage tree (this is a *consistency result* when tree error is defined as the number of false negative branches). Lastly, we consider the case of triplets, showing that there can be anomalous triplets when the probability of false positive errors is greater than zero. Our result suggests that quartets may be more robust to error than triplets when reconstructing cell lineage trees.

Our results also generalize beyond the `IS` model to any model *ℳ* of 2-state character evolution for which there are no anomalous quartets or triplets. An example of such a model is the infinite sites plus neutral Wright-Fisher (`IS+nWF`) model (Fisher 1923; Wright 1931) and its approximations (Hudson 2002). The `IS+nWF` model is a forward-time version of the MSC model. Mutations follow the IS assumption but evolve within a species tree, so deviations from a perfect phylogeny are due to genetic drift. Nevertheless, there are no anomalous triplets (see Supplementary Materials of Kuritzin et al. 2016) and or quartets (Theorem 1 in Molloy et al. 2021; also see Mendes and Hahn 2017) under the `IS+nWF` model. This led Springer and Gatesy (2016) use quartet-based methods to estimate species trees from retroelement insertion presence/absence patterns, and Molloy et al. (2021) showed this approach is consistent under the `IS+nWF` model. Since then, quartet-based methods have been applied to retroelement data sets for placental mammals (Doronina et al. 2022), bats (Korstian et al. 2022), and birds (Gatesy and Springer 2022).

Missing values are prevalent in these data sets; however, little was known about the performance of quartet-based methods when the input data are imperfect. A consequence of our study is that quartet-based methods, like ASTRAL, are consistent under the `IS+nWF` model, even when unbiased errors and missing values are introduced. This statement follows from combining Theorem 1 in Molloy et al. (2021) with Corollaries 1 and 3. Thus, our work addresses an open question from Molloy et al. (2021) and gives a positive result for recent systematic studies leveraging quartet-based methods on retroelement insertion presence/absence patterns. On the other hand, unbiased error is appropriate for modeling sequencing error, but error could be biased towards particular species or genes when calling retroelement insertions. Future work should investigate this issue further, looking at error and missingness biased towards particular species or (orthologous) positions of the genome. Lastly, although species trees are typically assumed to be binary, there could be hard polytomies, in which case the model tree would be non-binary. Our results for consistency with error defined as the number of false negatives (Lemma 6 and Corollary 3) extend to the MSC model, suggesting the utility of quartet-based methods in this setting with hard polytomies.

Coming back to tumor phylogenetics, our main result (Theorem 3) suggests the potential of quartet-based methods for reconstructing highly unresolved cell lineage trees, provided that the tree can be rooted and that false positive branches in the output tree can be effectively handled. The former is doable because the tree can be rooted on the edge incident to the healthy cell with no mutations. The latter is related to mapping mutations onto branches in the cell lineage tree (see Kozlov et al. 2022) as well as identifying which cells are members of the same clone or subclone (see Zafar et al. 2019). These tasks may need to be performed when using likelihood-based methods designed for cell lineage tree reconstruction, as such methods also return binary trees. Examples include ScisTree (Wu 2019), SiFit (Zafar et al. 2017), and CellPhy (Kozlov et al. 2022) (note that of these methods ScisTree makes the IS assumption but the other two do not).

In general, likelihood-based methods require explicitly estimating numeric parameters, like *α* and *β*, as well as exploring the space of cell lineage trees, which grows exponentially in the numbers of cells. In contrast, quartet-based approaches allow for error implicitly (without explicit estimation of *α* and *β*) and are often based on algorithmic techniques, like divide-and-conquer, that are quite fast in practice. That being said, quartet-based methods have been designed for species phylogenetics, where the number of leaves (species) is typically much less than the number of gene trees or characters. In tumor phylogenetics, the number of leaves (cells) can be much greater than than characters. This will likely to have consequences for runtime and accuracy (just consider that our consistency guarantees assume an infinite number of characters). Corollary 3 sheds light on a potential issue when using ASTRAL, that is, the construction of the constrained solution space σ may not be very successful for mutation data, especially if the number of mutations is small. However, there are other high quality heuristics for MQSS, including wQMC (Snir and Rao 2010, 2012; Avni et al. 2014) and wQFM (Reaz et al. 2014; Mahbub et al. 2021). wQFM, in particular, was recently shown to outperform ASTRAL in simulations (Mahbub et al. 2021). Moreover, even when the number of characters is small compared to the number of cells, the underlying model tree is likely to be highly unresolved. In this case, sampling different cells around the same branch may be a means of providing more data for estimation (this observation has already been leveraged by Kizilkale et al. 2022).

Lastly, we note that quartet-based methods, as presented here, fail to address doublets and copy number aberrations (CNAs), which also challenge cell lineage tree reconstruction. A doublet is a sequencing artifact where data provided for a single cell is really a mixture of two cells. This “hybrid” cell challenges the notion of tree-like evolution, motivating the development of methods for correcting doublets (Weber et al. 2021). If doublets can be effectively corrected, then their impact on quartet-based methods would be minimal. Alternatively, quartets may be useful for detecting doublets.

CNAs include duplications and losses of large sections of chromosomes (see review on methods for detecting CNAs by Mallory et al. 2020). CNA losses, in particular, have motivated the development of many new methods for reconstructing tumor phylogenies (El-Kebir 2018; Malikic et al. 2019; Satas et al. 2020). Some of these methods view CNA losses as false negatives (although these false negatives will be biased towards particular cells and mutations). In contrast, SCARLET (Satas et al. 2020) reconstructs a CNA tree and then uses it to constrain phylogeny reconstruction with the mutation data. Constraints have also been leveraged in species phylogenetics, including with ASTRAL (Rabiee and Mirarab 2020). Thus, the output of quartet-based methods could similarly be forced to obey the constraints of a CNA tree. To summarize, it seems possible that quartet-based methods can be tailored to address practical challenges in tumor phylogenetics, and we are optimistic about their potential for cell lineage tree reconstruction.

## 7. Acknowledgements

This work was conducted in preparation for the talk “Models and methods for reconstructing population-level evolutionary histories and relationships to tumor phylogenetics”, given at the U.S. National Cancer Institute’s Spring School in Algorithmic Cancer Biology on March 15, 2023. EKM thanks the organizers (M. El Kebir, S. Malikic, T.M. Przytycka, B. Raphael, S.C. Sahinalp, and M. Singh) for the invitation and the audience for their questions and engagement. The authors also thank Michael Nute for very helpful feedback on a preliminary version of this paper, especially our notation.

