## Supplementary Materials for "Quartets enable statistically consistent estimation of cell lineage trees under an unbiased error and missingness model"

Yunheng Han[1,2][0000−0003−0200−5924] and Erin K. Molloy[1,2,*][0000−0001−5553−3312]

[1] *Department of Computer Science, University of Maryland, College Park, USA*

[2] *University of Maryland Institute for Advanced Computer Studies, College Park, USA*

***

April 26, 2023

#### Contents

**List of Tables** **1**

**List of Figures** **1**

**1 Supplementary Proofs** **1**

**2 Supplementary Figures** **3**

**3 Supplementary Tables** **5**

#### List of Figures

S1 Triplet shapes . . . 4
S2 Triplet-based Phylogeny Estimation Pipeline . . . 4

### 1 Supplementary Proofs

**Lemma 1 from main text.** *Let* $0 \le \alpha < 1$ *and* $0 \le \beta < 1$. *Then,*

$$\big((1-\beta)^2(1-\alpha)^2 + \beta^2\alpha^2\big) - 2\beta(1-\beta)\alpha(1-\alpha) = \big(1-(\alpha+\beta)\big)^2 > 0 \tag{1}$$

*for* $\alpha + \beta \neq 1$. *If* $\alpha + \beta = 1$, *the inequality becomes an equality.*

*Proof.*

$$\begin{aligned}
(1-\beta)^2(1-\alpha)^2 + \beta^2\alpha^2 &> 2\beta(1-\beta)\alpha(1-\alpha)\\
(1-2\alpha+\alpha^2)(1-2\beta+\beta^2) + \alpha^2\beta^2 &> 2\alpha\beta(1-\alpha-\beta+\alpha\beta)\\
(1-2\beta+\beta^2-2\alpha+4\alpha\beta-2\alpha\beta^2+\alpha^2-2\alpha^2\beta+\alpha^2\beta^2)+\alpha^2\beta^2 &> 2\alpha\beta-2\alpha^2\beta-2\alpha\beta^2+2\alpha^2\beta^2\\
1-2\beta+\beta^2-2\alpha+2\alpha\beta+\alpha^2 &> 0\\
1+2\alpha\beta+\alpha^2+\beta^2-2(\alpha+\beta) &> 0\\
1+(\alpha+\beta)^2-2(\alpha+\beta) &> 0\\
\big(1-(\alpha+\beta)\big)^2 &> 0
\end{aligned}$$

□

**Lemma 3 from main text.** *There are no anomalous quartets under the* IS *model. Moreover, all quartet-informative patterns have zero probability except for one or both of the patterns corresponding to* $u(\sigma)$, *assuming* $u(\sigma)$ *is not a star.*

*Proof.* As discussed in the main text, it suffices to verify that there are no anomalous quartets for $\sigma$ with four leaves. We verify this statement by checking whether each of the six quartet-informative mutation patterns can be generated under the `IS` model given $\sigma$. We must consider that $\sigma$ can be any of the tree shapes shown in Figure 2 (from main text). Under the `IS` model, a mutation occurs on an edge $e = u \mapsto v$ and all leaves that are descendants of $v$ are in state 1 and all other leaves are in state 0; therefore, we only need to consider mutations occurring on the internal branches of each of the tree shapes (recall that mutations have a non-zero probability of occurring on any internal branch; otherwise the branch is contracted). Without loss of generality, we assume the tree shapes have their leaves labeled from left to right by label set $\{A, B, C, D\}$.

First, we consider the rooted tree shapes that when unrooted correspond to a quartet.

- For tree shape $\mathcal{S}_1$ with representative tree $((A, B), (C, D));$, a mutation on the left internal branch will produce pattern $ABCD = 1100$, and a mutation on the right internal branch will produce pattern 0011.
- For tree shape $\mathcal{S}_2$ with representative tree $(((A, B), C), D);$, a mutation on the internal branch closest to the leaves will produce pattern 1100 and a mutation on the other internal branch will produce pattern 1110.
- For tree shape $\mathcal{S}_3$ with representative tree $((A, B), C, D);$, a mutation on the only internal branch will produce pattern $ABCD = 1100$.

In all of these cases, the unrooted tree corresponds to $A, B|C, D$ and all quartet-informative pattern(s) correspond to $A, B|C, D$ (all other quartet-informative patterns have zero probability).

Second, we consider the rooted tree shapes that correspond to a star when unrooted.

- For tree shape $\mathcal{S}_4$ with representative tree $(A, B, C), D);$, a mutation on the only internal branch will produce pattern 1110.
- For tree shape $\mathcal{S}_5$ with representative tree $(A, B, C, D);$, there are no internal branches.

In all of these cases, the unrooted tree shape is a star and all quartet-informative patterns have zero probability.

By Definition 2 (from main text), there are no anomalous quartets under the `IS` model. □

**Lemma 5 from main text.** *There are no anomalous triplets under the* `IS` *model. Moreover, all triplet-informative patterns have zero probability except for the pattern corresponding to* $\sigma$*, assuming* $\sigma$ *is not non-binary.*

*Proof.* As discussed in the main text, it suffices to verify that there are no anomalous triplets for $\sigma$ with three leaves. We verify this statement by checking whether each of the three triplet-informative mutation patterns can be generated under the `IS` model given $\sigma$. We must consider that $\sigma$ can be any of the tree shapes shown in Figure S1. Under the `IS` model, a mutation occurs on an edge $e = u \mapsto v$ and all leaves that are descendants of $v$ are in state 1 and all other leaves are in state 0; therefore, we only need to consider mutations occurring on the internal branches of each of the tree shapes (recall that mutations have a non-zero probability of occurring on any internal branch; otherwise the branch is contracted). Without loss of generality, we assume the tree shapes have their leaves labeled from left to right by label set $\{A, B, C\}$.

- For tree shape $\mathcal{S}_6$ with representative tree $((A, B), C);$, a mutation on the only internal branch will produce pattern $ABC = 110$.

- For tree shape $\mathcal{S}_7$ with representative tree $(A, B, C);$, there are no internal branches.

In the first case, the rooted tree corresponds to $A, B|C$ and the only triplet-informative pattern also corresponds to $A, B|C$ (all other triplet-informative patterns have zero probability). In the second case, $\sigma$ is non-binary and all triplet-informative patterns have zero probability. By Definition 3 (from main text), there are no anomalous triplets under the `IS` model. □

### 2 Supplementary Figures

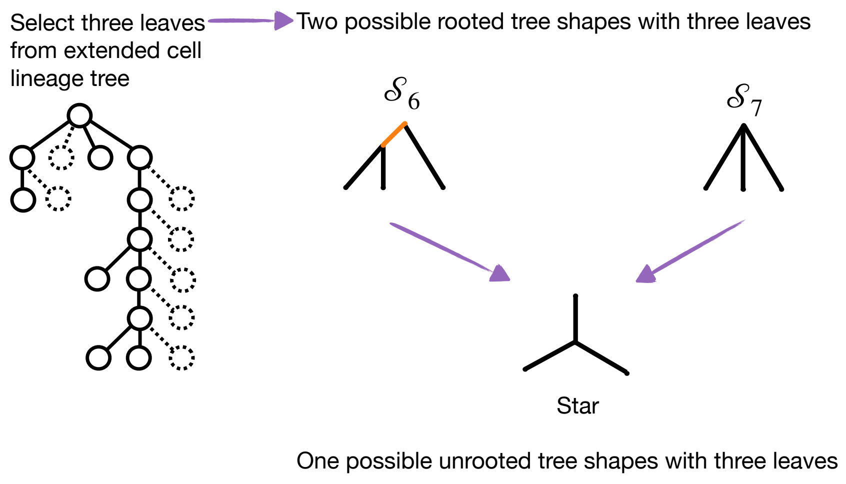

Figure S1: The two possible cell lineage tree shapes with three leaves. One of them ($\mathcal{S}_6$) has a non-trivial rooted topology (called a triplet), and the other ($\mathcal{S}_7$) has a trivial rooted topology. Both tree shapes $\mathcal{S}_6$ and $\mathcal{S}_7$ have a trivial topology when unrooted (called a star).

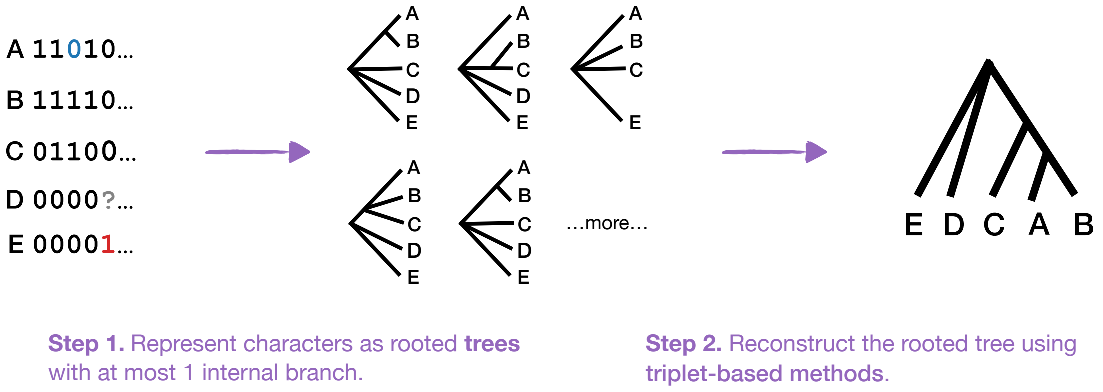

Figure S2: This figure shows a binary character matrix with one false negative (in blue), one false positive (in red), and one missing entry. The first step indicates how each character (column in the matrix) corresponds to a rooted tree with at most one internal branch under the IS assumption. The second step is to estimate the phylogeny by applying a heuristic for weighted triplet consensus. This approach is motivated by there being no anomalous triplets under the `IS+UEM` model for $\alpha = 0$ (Theorem 2 from the main text).

### 3 Supplementary Tables

| # of FP | # of FN | $\mathbb{P}_{\texttt{UE}}(D_{*,j}\|\alpha,\beta,G_{*,j})$ | $D_{*,j}$ derived from $G_{*,j}$ = **0000** | Quartet supported by $D_{*,j}$ |
|---|---|---|---|---|
| 0 | 0 | $(1-\alpha)^4$ | 0000 | none |
| 1 | 0 | $\alpha(1-\alpha)^3$ | 1000<br>0100<br>0010<br>0001 | none<br>none<br>none<br>none |
| 2 | 0 | $\alpha^2(1-\alpha)^2$ | 1100<br>1010<br>1001<br>0110<br>0101<br>0011 | $A,B\|C,D$<br>$A,C\|B,D$<br>$A,D\|B,C$<br>$A,D\|B,C$<br>$A,C\|B,D$<br>$A,B\|C,D$ |
| 3 | 0 | $\alpha^3(1-\alpha)$ | 1110<br>1101<br>1011<br>0111 | none<br>none<br>none<br>none |
| 4 | 0 | $\alpha^4$ | 1111 | none |

Table S1: Above we show the characters that can be generated from **character #0** ($ABCD = 0000$) under the `UE` model. Recall that false positives ($0 \to 1$; shown in red) occur with probability $\alpha$, false negatives ($1 \to 0$; shown in blue) occur with probability $\beta$, and these errors are independent and identically distributed across mutations and cells.

| # of FP | # of FN | $\mathbb{P}_{\texttt{UE}}(D_{*,j}\|\alpha,\beta,G_{*,j})$ | $D_{*,j}$ derived from $G_{:*,j}$ = **1111** | Quartet supported by $D_{*,j}$ |
|---|---|---|---|---|
| 0 | 0 | $(1-\beta)^4$ | 1111 | none |
| 0 | 1 | $\beta(1-\beta)^3$ | 0111<br>1011<br>1101<br>1110 | none<br>none<br>none<br>none |
| 0 | 2 | $\beta^2(1-\beta)^2$ | 0011<br>0101<br>0110<br>1001<br>1010<br>1100 | $A,B\|C,D$<br>$A,C\|B,D$<br>$A,D\|B,C$<br>$A,D\|B,C$<br>$A,C\|B,D$<br>$A,B\|C,D$ |
| 0 | 3 | $\beta^3(1-\beta)$ | 0001<br>0010<br>0100<br>1000 | none<br>none<br>none<br>none |
| 0 | 4 | $\beta^4$ | 0000 | none |

Table S2: Above we show the 4-characters that can be generated from **4-character #15** ($ABCD = 1111$) under the `UE` model.

| # of FP | # of FN | $\mathbb{P}_{\texttt{UE}}(D_{*,j}\|\alpha,\beta,G_{*,j})$ | $D_{*,j}$ derived from $G_{*,j}$ = **0001** | $G_{*,j}$ = **0100** | $G_{*,j}$ = **0100** | $G_{*,j}$ = **1000** | Quartet supported by $D_{*,j}$ |
|---|---|---|---|---|---|---|---|
| 0 | 0 | $(1-\alpha)^3(1-\beta)$ | 0001 | 0010 | 0100 | 1000 | none |
| 0 | 1 | $(1-\alpha)^3\beta$ | 0000 | 0000 | 0000 | 0000 | none |
| 1 | 0 | $\alpha(1-\alpha)^2(1-\beta)$ | 1001<br>0101<br>0011 | 0110<br>1010<br>0011 | 0110<br>0101<br>1100 | 1001<br>1010<br>1100 | $A,D\|B,C$<br>$A,C\|B,D$<br>$A,B\|B,C$ |
| 1 | 1 | $\alpha(1-\alpha)^2\beta$ | 1000<br>0100<br>0010 | 0100<br>1000<br>0001 | 0010<br>0001<br>1000 | 0001<br>0010<br>0100 | none<br>none<br>none |
| 2 | 0 | $\alpha^2(1-\alpha)(1-\beta)$ | 1101<br>1011<br>0111 | 1110<br>1011<br>0111 | 1110<br>1101<br>0111 | 1110<br>1101<br>1011 | none<br>none<br>none |
| 2 | 1 | $\alpha^2(1-\alpha)\beta$ | 1100<br>1010<br>0110 | 1100<br>0101<br>1001 | 0011<br>1010<br>1001 | 0011<br>0101<br>0110 | $A,B\|C,D$<br>$A,C\|B,D$<br>$A,D\|B,C$ |
| 3 | 0 | $\alpha^3(1-\beta)$ | 1111 | 1111 | 1111 | 1111 | none |
| 3 | 1 | $\alpha^3\beta$ | 1110 | 1101 | 1011 | 0111 | none |

Table S3: Above we show the characters that can be generated from **character #1** ($ABCD$ = 1000), **character #2** (0100), **character #4** (0010), and **character #8** (0001) under the UE model.

| # of FP | # of FN | $\mathbb{P}_{\texttt{UE}}(D_{*,j}\|\alpha,\beta,G_{*,j})$ | $D_{:,j}$ derived from $G_{*,j}$ = **0111** | $G_{*,j}$ = **1011** | $G_{*,j}$ = **1101** | $G_{*,j}$ = **1110** | Quartet supported by $D_{*,j}$ |
|---|---|---|---|---|---|---|---|
| 0 | 0 | $(1-\beta)^3(1-\alpha)$ | 0111 | 1011 | 1101 | 1110 | none |
| 0 | 1 | $\beta(1-\beta)^2(1-\alpha)$ | 0110<br>0101<br>0011 | 1001<br>1010<br>0011 | 1001<br>0101<br>1100 | 0110<br>1010<br>1100 | $A,D\|B,C$<br>$A,C\|B,D$<br>$A,B\|C,D$ |
| 0 | 2 | $\beta^2(1-\beta)(1-\alpha)$ | 0001<br>0010<br>0100 | 0001<br>0010<br>1000 | 0001<br>0100<br>1000 | 0010<br>0100<br>1000 | none<br>none<br>none |
| 0 | 3 | $\beta^3(1-\alpha)$ | 0000 | 0000 | 0000 | 0000 | none |
| 1 | 0 | $(1-\beta)^3\alpha$ | 1111 | 1111 | 1111 | 1111 | none |
| 1 | 1 | $\beta(1-\beta)^2\alpha$ | 1011<br>1111<br>1110 | 0111<br>1101<br>1110 | 0111<br>1011<br>1110 | 0111<br>1011<br>1101 | none<br>none<br>none |
| 1 | 2 | $\beta^2(1-\beta)\alpha$ | 1100<br>1010<br>1001 | 1100<br>0101<br>0110 | 0011<br>1010<br>0110 | 0011<br>0101<br>1001 | $A,B\|C,D$<br>$A,C\|B,D$<br>$A,D\|B,C$ |
| 1 | 3 | $\beta^3\alpha$ | 1000 | 0100 | 0010 | 0001 | none |

Table S4: Above we show the 4-characters that can be generated from **4-character #7** ($ABCD$ = 0111), **4-character #11** ($ABCD$ = 1011), **4-character #13** ($ABCD$ = 1101), and **4-character #14** ($ABCD$ = 1110) under the UE model.

| # of FP | # of FN | $\mathbb{P}_{\texttt{UE}}(D_{*,j}\|\alpha,\beta,G_{*,j})$ | $D_{*,j}$ derived from $G_{*,j}$ = **0101** | $G_{*,j}$ = **1010** | Quartet supported by $D_{*,j}$ |
|---|---|---|---|---|---|
| 0 | 0 | $(1-\alpha)^2(1-\beta)^2$ | 0101 | 1010 | $A,C\|B,D$ |
| 0 | 1 | $\beta(1-\beta)(1-\alpha)^2$ | 0001<br>0100 | 0010<br>1000 | none<br>none |
| 0 | 2 | $\beta^2(1-\beta)^2$ | 0000 | 0000 | none |
| 1 | 0 | $(1-\beta)^2\alpha(1-\alpha)$ | 1101<br>0111 | 1110<br>1011 | none<br>none |
| 1 | 1 | $\beta(1-\beta)\alpha(1-\alpha)$ | 1001<br>1100<br>0011<br>0110 | 0110<br>0011<br>1100<br>1001 | $A,D\|B,C$<br>$A,B\|C,D$<br>$A,B\|C,D$<br>$A,D\|B,C$ |
| 1 | 2 | $\beta^2\alpha(1-\alpha)$ | 1000<br>0010 | 0100<br>0001 | none<br>none |
| 2 | 0 | $(1-\beta)^2\alpha^2$ | 1111 | 1111 | none |
| 2 | 1 | $\beta(1-\beta)\alpha^2$ | 1011<br>1110 | 0111<br>1101 | none<br>none |
| 2 | 2 | $\beta^2\alpha^2$ | 1010 | 0101 | $A,C\|B,D$ |

Table S5: Above we show the 4-characters that can be generated from **4-character #5** ($ABCD = 0101$) and **4-character #10** (1010) under the UE model.

| # of FP | # of FN | $\mathbb{P}_{\texttt{UE}}(D_{*,j}\|\alpha,\beta,G_{*,j})$ | $D_{*,j}$ derived from $G_{*,j}$ = **0110** | $G_{*,j}$ = **1001** | Quartet supported by $D_{*,j}$ |
|---|---|---|---|---|---|
| 0 | 0 | $(1-\alpha)^2(1-\beta)^2$ | 0110 | 1001 | $A,D\|B,C$ |
| 0 | 1 | $\beta(1-\beta)(1-\alpha)^2$ | 0010<br>0100 | 0001<br>1000 | none<br>none |
| 0 | 2 | $\beta^2(1-\beta)^2$ | 0000 | 0000 | none |
| 1 | 0 | $(1-\beta)^2\alpha(1-\alpha)$ | 1110<br>0111 | 1101<br>1011 | none<br>none |
| 1 | 1 | $\beta(1-\beta)\alpha(1-\alpha)$ | 1010<br>1100<br>0011<br>0101 | 0101<br>0011<br>1100<br>1010 | $A,C\|B,D$<br>$A,B\|C,D$<br>$A,B\|C,D$<br>$A,C\|B,D$ |
| 1 | 2 | $\beta^2\alpha(1-\alpha)$ | 1000<br>0001 | 0100<br>0010 | none<br>none |
| 2 | 0 | $(1-\beta)^2\alpha^2$ | 1111 | 1111 | none |
| 2 | 1 | $\beta(1-\beta)\alpha^2$ | 1011<br>1101 | 0111<br>1110 | none<br>none |
| 2 | 2 | $\beta^2\alpha^2$ | 1001 | 0110 | $A,D\|B,C$ |

Table S6: Above we show the 4-characters that can be generated from **4-character #6** ($ABCD = 0110$) and **4-character #9** (1001) under the UE model.

| # of FP | # of FN | $\mathbb{P}_{\texttt{UE}}(D_{*,j} \mid \alpha, \beta, G_{*,j})$ | $D_{*,j}$ derived from $G_{*,j}$ = **000** | Triplet supported by $D_{*,j}$ |
|---|---|---|---|---|
| 0 | 0 | $(1-\alpha)^3$ | 000 | none |
| 1 | 0 | $\alpha(1-\alpha)^2$ | 100<br>010<br>001 | none<br>none<br>none |
| 2 | 0 | $\alpha^2(1-\alpha)$ | 110<br>101<br>011 | $C \mid A, B$<br>$B \mid A, C$<br>$A \mid B, C$ |
| 3 | 0 | $\alpha^3$ | 111 | none |

Table S7: Above we show the 3-characters that can be generated from **3-character #0** ($ABC = 000$) under the UE model. Recall that false positives ($0 \to 1$; shown in red) occur with probability $\alpha$, false negatives ($1 \to 0$; shown in blue) occur with probability $\beta$, and these errors are independent and identically distributed across mutations and cells.

| # of FP | # of FN | $\mathbb{P}_{\texttt{UE}}(D_{*,j} \mid \alpha, \beta, G_{*,j})$ | $D_{*,j}$ derived from $G_{*,j}$ = **111** | Triplet supported by $D_{*,j}$ |
|---|---|---|---|---|
| 0 | 0 | $(1-\beta)^3$ | 111 | none |
| 0 | 1 | $\beta(1-\beta)^2$ | 011<br>101<br>110 | $A \mid B, C$<br>$B \mid A, C$<br>$C \mid A, B$ |
| 0 | 2 | $\beta^2(1-\beta)$ | 001<br>010<br>100 | none<br>none<br>none |
| 0 | 3 | $\beta^3$ | 000 | none |

Table S8: Above we show the 3-characters that can be generated from **3-character #7** ($ABC = 111$) under the UE model.

| # of FP | # of FN | $\mathbb{P}_{\texttt{UE}}(D_{*,j} \mid \alpha, \beta, G_{*,j})$ | $D_{*,j}$ derived from $G_{*,j}$ = **001** | Triplet supported by $D_{*,j}$ |
|---|---|---|---|---|
| 0 | 0 | $(1-\alpha)^2(1-\beta)$ | 001 | none |
| 1 | 0 | $\alpha(1-\alpha)(1-\beta)$ | 101<br>011 | $B \mid A, C$<br>$A \mid B, C$ |
| 2 | 0 | $\alpha^2(1-\beta)$ | 111 | none |
| 0 | 1 | $(1-\alpha)^2\beta$ | 000 | none |
| 1 | 1 | $\alpha(1-\alpha)\beta$ | 100<br>010 | none<br>none |
| 2 | 1 | $\alpha^2\beta$ | 110 | $C \mid A, B$ |

Table S9: Above we show the 3-characters that can be generated from **3-character #1** ($ABC = 001$) under the UE model.

| # of FP | # of FN | $\mathbb{P}_{\texttt{UE}}(D_{*,j} \mid \alpha, \beta, G_{*,j})$ | $D_{*,j}$ derived from $G_{*,j}$ = **010** | Triplet supported by $D_{*,j}$ |
|---|---|---|---|---|
| 0 | 0 | $(1-\alpha)^2(1-\beta)$ | 010 | none |
| 1 | 0 | $\alpha(1-\alpha)(1-\beta)$ | 110<br>011 | $C\|A, B$<br>$A\|B, C$ |
| 2 | 0 | $\alpha^2(1-\beta)$ | 111 | none |
| 0 | 1 | $(1-\alpha)^2\beta$ | 000 | none |
| 1 | 1 | $\alpha(1-\alpha)\beta$ | 100<br>001 | none<br>none |
| 2 | 1 | $\alpha^2\beta$ | 101 | $B\|A, C$ |

Table S10: Above we show the 3-characters that can be generated from **3-character #2** ($ABC = 010$) under the UE model.

| # of FP | # of FN | $\mathbb{P}_{\texttt{UE}}(D_{*,j} \mid \alpha, \beta, G_{*,j})$ | $D_{*,j}$ derived from $G_{*,j}$ = **100** | Triplet supported by $D_{*,j}$ |
|---|---|---|---|---|
| 0 | 0 | $(1-\alpha)^2(1-\beta)$ | 100 | none |
| 1 | 0 | $\alpha(1-\alpha)(1-\beta)$ | 110<br>101 | $C\|A, B$<br>$B\|A, C$ |
| 2 | 0 | $\alpha^2(1-\beta)$ | 111 | none |
| 0 | 1 | $(1-\alpha)^2\beta$ | 000 | none |
| 1 | 1 | $\alpha(1-\alpha)\beta$ | 010<br>001 | none<br>none |
| 2 | 1 | $\alpha^2\beta$ | 011 | $A\|B, C$ |

Table S11: Above we show the 3-characters that can be generated from **3-character #4** ($ABC = 100$) under the UE model.

| # of FP | # of FN | $\mathbb{P}_{\texttt{UE}}(D_{*,j} \mid \alpha, \beta, G_{*,j})$ | $D_{*,j}$ derived from $G_{*,j}$ = **011** | Triplet supported by $D_{*,j}$ |
|---|---|---|---|---|
| 0 | 0 | $(1-\alpha)(1-\beta)^2$ | 011 | $A\|B, C$ |
| 1 | 0 | $\alpha(1-\beta)^2$ | 111 | none |
| 0 | 1 | $(1-\alpha)\beta(1-\beta)$ | 001<br>010 | none<br>none |
| 0 | 2 | $(1-\alpha)\beta^2$ | 000 | none |
| 1 | 1 | $\alpha\beta(1-\beta)$ | 101<br>110 | $B\|A, C$<br>$C\|A, B$ |
| 1 | 2 | $\alpha\beta^2$ | 100 | none |

Table S12: Above we show the 3-characters that can be generated from **3-character #3** ($ABC = 011$) under the UE model.

| # of FP | # of FN | $\mathbb{P}_{\texttt{UE}}(D_{*,j}\mid\alpha,\beta,G_{*,j})$ | $D_{*,j}$ derived from $G_{*,j}=\mathbf{101}$ | Triplet supported by $D_{*,j}$ |
|---|---|---|---|---|
| 0 | 0 | $(1-\alpha)(1-\beta)^2$ | 101 | $B\|A,C$ |
| 0 | 1 | $(1-\alpha)\beta(1-\beta)$ | 001<br>100 | none<br>none |
| 0 | 2 | $(1-\alpha)\beta^2$ | 000 | none |
| 1 | 0 | $\alpha(1-\beta)^2$ | 111 | none |
| 1 | 1 | $\alpha\beta(1-\beta)$ | 011<br>110 | $A\|B,C$<br>$C\|A,B$ |
| 1 | 2 | $\alpha\beta^2$ | 010 | none |

Table S13: Above we show the 3-characters that can be generated from **3-character #5** ($ABC = 101$) under the `UE` model.

| # of FP | # of FN | $\mathbb{P}_{\texttt{UE}}(D_{*,j}\mid\alpha,\beta,G_{*,j})$ | $D_{*,j}$ derived from $G_{*,j}=\mathbf{110}$ | Triplet supported by $D_{*,j}$ |
|---|---|---|---|---|
| 0 | 0 | $(1-\alpha)(1-\beta)^2$ | 110 | $C\|A,B$ |
| 0 | 1 | $(1-\alpha)\beta(1-\beta)$ | 010<br>100 | none<br>none |
| 0 | 2 | $(1-\alpha)\beta^2$ | 000 | none |
| 1 | 0 | $\alpha(1-\beta)^2$ | 111 | none |
| 1 | 1 | $\alpha\beta(1-\beta)$ | 011<br>101 | $A\|B,C$<br>$B\|A,C$ |
| 1 | 2 | $\alpha\beta^2$ | 001 | none |

Table S14: Above we show the 3-characters that can be generated from **3-character #6** ($ABC = 110$) under the `UE` model.

| ID | Character $G_{*,j}$ $ABCD$ | Quartet supported by $G_{*,j}$ | $\mathcal{S}_1$ $((A,B),(C,D));$ | $\mathcal{S}_2$ $(((A,B),C),D);$ | $\mathcal{S}_3$ $((A,B),C,D);$ | $\mathcal{S}_4$ $((A,B,C),D);$ | $\mathcal{S}_5$ $(A,B,C,D);$ |
|---|---|---|---|---|---|---|---|
| 0 | 0000 | none | $\geq 0$ | $\geq 0$ | $\geq 0$ | $\geq 0$ | $\geq 0$ |
| 15 | 1111 | none | $\geq 0$ | $\geq 0$ | $\geq 0$ | $\geq 0$ | $\geq 0$ |
| 7 | 0111 | none | ✗ | ✗ | ✗ | ✗ | ✗ |
| 8 | 1000 | none | $\geq 0$ | $\geq 0$ | $\geq 0$ | $\geq 0$ | $\geq 0$ |
| 4 | 0100 | none | $\geq 0$ | $\geq 0$ | $\geq 0$ | $\geq 0$ | $\geq 0$ |
| 11 | 1011 | none | ✗ | ✗ | ✗ | ✗ | ✗ |
| 2 | 0010 | none | $\geq 0$ | $\geq 0$ | $\geq 0$ | $\geq 0$ | $\geq 0$ |
| 13 | 1101 | none | ✗ | ✗ | ✗ | ✗ | ✗ |
| 1 | 0001 | none | $\geq 0$ | $\geq 0$ | $\geq 0$ | $\geq 0$ | $\geq 0$ |
| 14 | 1110 | none | ✗ | ✓ | ✗ | ✓ | ✗ |
| 3 | 0011 | $A,B\|C,D$ | ✓ | ✗ | ✗ | ✗ | ✗ |
| 12 | 1100 | $A,B\|C,D$ | ✓ | ✓ | ✓ | ✗ | ✗ |
| 5 | 0101 | $A,C\|B,D$ | ✗ | ✗ | ✗ | ✗ | ✗ |
| 10 | 1010 | $A,C\|B,D$ | ✗ | ✗ | ✗ | ✗ | ✗ |
| 6 | 0110 | $A,D\|B,C$ | ✗ | ✗ | ✗ | ✗ | ✗ |
| 9 | 1001 | $A,D\|B,C$ | ✗ | ✗ | ✗ | ✗ | ✗ |

Table S15: The table above shows the probability of each of the 16 possible binary characters on four leaves for each of the 5 possible tree shapes (using the representative leaf labeling indicated by the newick string). ✓ indicates the character has strictly greater than 0 probability under the **infinite sites model** given the model tree $\sigma$ (Figure 2 in main text); ✗ indicates the character has 0 probability; $\geq 0$ indicates the probability has either greater than or equal to zero. The characters $1000, 0100, 0010, 0001$ can have zero probability in the case of fake nodes (because there cannot be mutations on the edges incident to fake nodes, which represent ancestral cells). The character 1111 can have zero probability in the case that one of the cells sampled is the fake root (as this cell will not carry any mutations); it could have probability greater than zero if we sample four cells that all carry the mutation. The character 0000 can have probability greater than zero because we could sample four cells that all do not carry the mutation.

| ID | Character $G_{*,j}$ $ABC$ | Triplet supported by $G_{*,j}$ | $\mathcal{S}_6$ $((A,B),C);$ | $\mathcal{S}_7$ $(A,B,C);$ |
|---|---|---|---|---|
| 0 | 000 | none | $\geq 0$ | $\geq 0$ |
| 7 | 111 | none | $\geq 0$ | $\geq 0$ |
| 1 | 001 | none | $\geq 0$ | $\geq 0$ |
| 6 | 110 | $((A,B),C);$ | ✓ | ✗ |
| 2 | 010 | none | $\geq 0$ | $\geq 0$ |
| 5 | 101 | $((A,C),B);$ | ✗ | ✗ |
| 2 | 100 | none | $\geq 0$ | $\geq 0$ |
| 5 | 011 | $((B,C),A);$ | ✗ | ✗ |

Table S16: The table above shows the probability of each of the 8 possible binary characters on three leaves for each of the 2 possible tree shapes (using the representative leaf labeling indicated by the newick string). ✓ indicates the character has strictly greater than 0 probability under the **infinite sites model** given the model tree $\sigma$ (Figure S1); ✗ indicates the character has 0 probability; $\geq 0$ indicates the probability has either greater than or equal to zero. The characters $100, 010, 001$ can have zero probability in the case of fake nodes (because there cannot be mutations on the edges incident to fake nodes, which represent ancestral cells). The character 111 can have zero probability in the case that one of the cells sampled is the fake root (as this cell will not carry any mutations); it could have probability greater than zero if we sample four cells that all carry the mutation. The character 000 can have probability greater than zero because we could sample four cells that all do not carry the mutation.
